# Characterization of a High-Affinity Copper Transporter in the White-Nose Syndrome Causing Fungal Pathogen *Pseudogymnoascus destructans*

**DOI:** 10.1101/2024.08.28.610057

**Authors:** Alyssa D. Friudenberg, Saika Anne, Ryan L. Peterson

## Abstract

Copper is an essential micronutrient and the ability to scavenge tightly bound or trace levels of copper ions at the host-pathogen interface is vital for fungal proliferation in animal hosts. Recent studies suggest that trace metal ion acquisition is critical for the establishment and propagation of *Pseudogymnoascus destructans*, the fungal pathogen responsible for white-nose syndrome (WNS), on their bat host. However, little is known about these metal acquisition pathways in *P. destructans*. In this study, we report the characterization of the *P. destructans* high-affinity copper transporter VC83_00191 (*Pd*CTR1a), which is implicated as a virulence factor associated with the WNS disease state. Using *Saccharomyces cerevisiae* as a recombinant expression host, we find that *Pd*CTR1a localizes to the cell surface plasma membrane and can efficiently traffic Cu-ions into the yeast cytoplasm. Complementary studies in the native *P. destructans* fungus provide evidence that *Pd*CTR1a transcripts and protein levels are dictated by Cu-bioavailability in the growth media. Our study demonstrates that *Pd*CTR1a is a functional high-affinity copper transporter and is relevant to Cu-homeostasis pathways in *P. destructans*.

## Introduction

The psychrophilic fungus *Pseudogymnoascus destructans* is responsible for white-nose syndrome (WNS) disease which causes burn-like infection lesions on the wings and muzzle of bats.^1-3^ This infectious fungal disease is prevalent in North America and affects many cave-dwelling bat species.^4^ Since WNS was first detected in a New York cave ecosystem in 2006, *P. destructans* has spread across North America and can be found in 40 US States and 9 Canadian provinces.^5, 6^ WNS has killed millions of North American bats and can cause up to 100% mortality in some bat populations.^7^ Currently, 12 bat species have been confirmed with WNS, including the gray bat, northern long-eared bat, Indiana bat, and tricolor bat, which are endangered North American bat species.^5^ There is no approved treatment for WNS. However, there are ongoing efforts that aim to slow *P. destructans* transmission rates and to reduce the fungal burden on infected hibernating bats to help preserve North American bat biodiversity.

*Pseudogymnoascus destructans* spores are resilient and *P. destructans* is known to utilize a wide range of nitrogen and carbon sources to propagate efficiently in diverse environmental conditions including the soil, cave hibernacula, and susceptible bat hosts.^8-10^ Recent studies support the hypothesis that *P. destructans* may have originated as a soil-dwelling plant-infecting pathogen that adapted to thrive on bat animal hosts.^11^ Thus, *P. destructans* may be an example of an “accidental pathogen” that is able to utilize its evolved growth mechanisms to thrive on susceptible bat hosts.^12^ Supporting this hypothesis is the observation that white-nose syndrome disease does not infect all bats equally and the transcript levels of *P. destructans* virulence factors correlate to fungal biomass.^13^ This may indicate that genes naturally supporting the saprophytic lifestyle of *P. destructans* may have a dual role in establishing and propagating on the animal host.

The ability of microbial fungal pathogens to secure trace metal micronutrients such as copper are essential to maintaining cellular redox homeostasis needed for efficient propagation.^14-17^ Animal hosts have the ability to restrict access to essential metal ions, known as the process of nutritional immunity, which can involve the mobilization of metals to other tissues or the secretion of metal-binding proteins such as calprotectin and the S100-family of proteins.^18-20^ Transcriptomics studies into the host transcriptional response at active fungal infection sites on WNS-positive bats detect elevated levels of S100 transcripts,^21^ suggesting that the bat-host may be employing a metal-withholding strategy to slow *P. destructans* propagation on infected tissue. Complementary *P. destructans* transcriptomics studies have revealed that several high-affinity metal transporter genes are associated with WNS fungal pathology.^22^ Together, these transcriptomics studies in addition to the chemical identification of siderophores^23^ and riboflavin^24^ at *P. destructans* infection sites indicate that the environmental niche at the host-pathogen interface is devoid of essential transition metal micronutrients.

The ability of pathogenic fungi to scavenge essential copper micronutrients from their host is important for fungal propagation and fungal invasion across host tissues. This copper is needed to supply essential copper cofactors to metalloenzyme virulence factors such as Cu-Superoxide dismutases (SODs)^25^ and laccases^26^. Fungal pathogens can utilize two primary pathways to scavenge essential copper including the high-affinity copper transporters (CTRs) or isocyanide calkophore small molecules.^27, 28^ The family of copper transporters (CTRs) is the most conserved and well-studied class of Cu-scavenging proteins in the fungal kingdom and are essential virulence factors in many human-infecting fungal pathogens.^29, 30^ Recent transcriptomics studies at WNS disease sites have implicated the putative CTR transporter, VC83_00191, as an essential virulence factor in *P. destructans*.^22^

In this work, we identify *P. destructans* VC83_00191, designated as *Pd*CTR1a, as a functional high-affinity copper transporter that participates in the *P. destructans* Cu-stress response. Using baker’s yeast *Saccharomyces cerevisiae*, we demonstrate that recombinant *Pd*CTR1a localizes to the plasma membrane and can efficiently traffic Cu-ions into the yeast cytoplasm. Complementary studies in the native *P. destructans* fungus indicate that *P. destructans Pd*CTR1a and the second CTR transporter VC83_04814, designated as *Pd*CTR1b are transcriptionally regulated by Cu-stress. Furthermore, we detect the presence of *Pd*CTR1a protein in *P. destructans* lysates grown under copper-restrictive growth conditions. These data present the first biochemical characterization of putative *P. destructans* transmembrane protein as virulence factors implicated in WNS-disease and the investigation of the copper cell biology *P. destructans*.

### 2.1 Strains and media

The *Saccharomyces cerevisiae* strains used in this study include the BY4741 (MATa his3Δ1 leu2Δ0 met15Δ0 ura3Δ0) and BY4741ctr1Δ (MATa his3Δ1 leu2Δ0 met15Δ0 ura3Δ0 ctr1:kanMX) from the Yeast Knockout collection (Horizon Discovery Group plc). *Pseudogymnoascus destructans*, formally *Geomyces destructans MYA-4855* (Strain ID 20631-21), was purchased from the American Tissue Culture Collection (ATCC). All fungi were propagated using YPD or chemically defined growth media.

Chemically defined Synthetic Complete growth media SC-His and SC-Ura consisted of the base formulation of 1.7 g/L Yeast Nitrogen Base (YNB), 5.0 g/L ammonium sulfate, 20 g/L glucose, 1.83 g/L SC -His-Ura (Sunrise Science Products; Knoxville, TN). One hundred milligrams per liter of uracil (TCI Chemicals; Portland, OR) or L-Histidine (RPI; Mt. Prospect, IL) was added to the SC-base formulation to generate SC-His or SC-Ura respective growth media. Solid-supported agarose SC-His and SC-Ura plates were made by adding 20 g/L agarose and 1 pellet (∼200 mg) of sodium hydroxide before autoclave sterilization. SC-His plates containing glycerol and ethanol carbon sources were made in a similar process with 20 g/L agarose, 1.83 g/L SC-His-Ura, 5.0 g/L ammonium sulfate, 100 mg/L uracil, 2% (v/v) ethanol, 2% (v/v) glycerol, and a pellet of sodium hydroxide. Rich YPD growth media consisted of 10 g/L yeast extract, 20 g/L peptone and 20 g/l glucose; 20 g/L of agarose was added prior to autoclave sterilization to make solid growth plates.

### 2.2 Bioinformatic Methods

Copper transporter (CTR) homologs from *Saccharomyces cerevisiae, Cryptococcus neoformans*, and *Pseudogymnoascus destructans* were retrieved from the NCBI Gene databank with the gene accession identifiers YPR124W, YHR175W, CNAG_00979, CNAG_01872, CNAG_07701 VC83_00191, and VC83_04814. Multiple sequence alignments were performed using Clustal omega.^31^ The server DeepTMHMM prediction server (Technical University of Denmark) was used to assign CTR homolog transmembrane domains.^32^

### 2.3 Recombinant plasmid construction

The generation of cytosolic cyGFP and *Pd*CTR1a-GFP expression cassettes driven by the *S. cerevisiae* Ctr1 promoter and terminator was achieved using a synthesized gene block purchased by Twist Bioscience (San Francisco, CA) that was placed in the BSA1-restriction sites of the holding vector pYTK-DN6 (Addgene.org).^33^ The native *S. cerevisiae* Ctr1 promoter consisted of the 610 bp sequence region prior to the Ctr1 start codon and Ctr1 terminator consisted of 510 bp sequence region after the Ctr1 stop codon. The gene block encoding for *Pseudogymnoascus destructans* CTR1a isoform was codon optimized for *S. cerevisiae* by Twist Bioscience and was engineered to include a dual Strep-tag II linker between the CTR1a and GFP proteins.

The GFP yeast expression plasmids pAF23 and pAF24 were generated using the PCR-amplified products containing engineered primers oRLP497 and oRLP498 with Spe1/EcoR1 compatible restriction site overhangs after BSA1 restriction enzyme digest. The resulting insert was incorporated into the unique EcoR1 and Spe1 sites in the multi-cloning site of pRS313. The complete sequence of plasmids pAF23 and pAF24 is available in the SI and these plasmids can be requested from Addgene.org (Watertown, MA).

### 2.4 Generation of recombinant yeast strains

The plasmids pRS313, pAF23, and pAF24 were transformed using the Li-Acetate PEG transformation protocol^34^ into *S. cerevisiae* BY4741 (WT) and BY4741*ctr1*Δ (*ctr1*Δ) yeast backgrounds referred to as WT and *ctr1*Δ strains here on forward. The six recombinant *S. cerevisiae* strains used in this study are overviewed in Table 1.

**Table 1.**
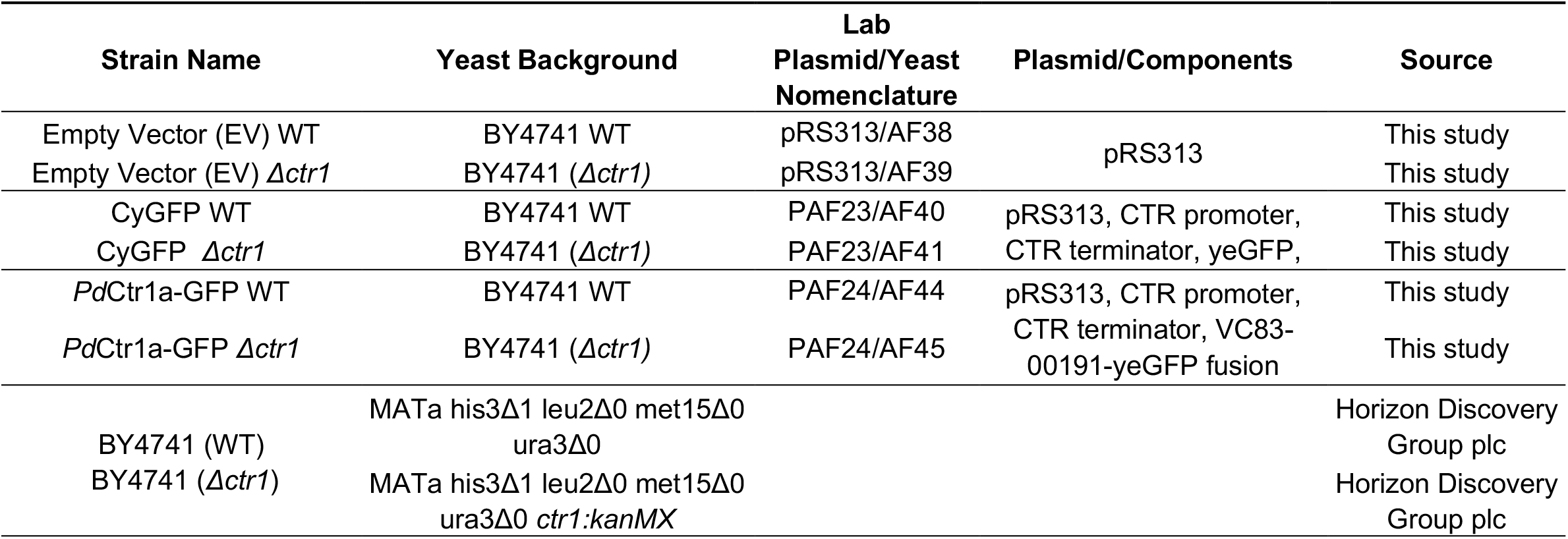
Table of Yeast strains used in this study.

### 2.5 Generation of PdCTR1a antiserum

Custom anti-serum products were achieved using the standard 90-New Zealand rabbit antibody services from Cocalico Biologicals, Inc. (Reamstown, PA; Assurance number D16-00398 (A3669-01)). The peptide fragments *KARQEARWLDCEMHRRYC* and *GKPGLRERVALHKDAKC* was selected as the *Pd*CTR1a antigen targets and coupled to Keyhole limpet hemocyanin (KLH) as the carrier protein. The co-injection of both *Pd*CTR1a peptide-KLH constructs was used for antibody production in rabbit hosts.

### 2.6 Flow cytometry assays

Yeast strains were cultivated in SC-His medium supplemented with 50 μM CuSO_4_ and 100 μM Bathocuproine disulfonate (BCS) overnight. The following morning, cells were harvested at an optical density of ∼0.8 at 600 nm (OD_600_) and subsequently diluted 1/10 in Milli-Q water prior to flow cytometry analysis. Cellular GFP fluorescence was quantified using a Beckman Coulter Cytoflex-S flow cytometer using the 488-nm blue laser line and 530/30-nm band-pass (GFP) filter. Data analysis was performed using Foreada (floreada.io); GFP fluorescence intensity was normalized to empty vector yeast given an intensity value of 10^3^ fluorescence units.

### 2.7 Dot blot assay

Dot blot experiments were performed using freshly prepared yeast cultures grown on SC-His plates at room temperature for 3 days. Cultured yeast were transferred from the growth plate to a microcentrifuge tube containing 1 mL of liquid SC-His media. The OD_600_ of this solution was acquired using the Cary 60 UV-Vis Spectrophotometer. Samples were normalized to yield 1 ml of a 1 OD_600_ stock solution in Milli-Q water. Subsequential 1:10 dilutions serial dilutions were made using Milli-Q water and 5 μL of each sample was deposited onto solid SC-His or SC-His with 2% glycerol and 2% ethanol + 5.0 μM supplemental copper sulfate growth plates. The resulting plates were incubated at 28 °C for 5-6 days. Images of the growth plates were captured using a digital camera.

### 2.8 Western blot analysis

Recombinant *S. cerevisiae* yeast transformants were cultivated in 10 mL of Sc-His media at 30°C to mid-log phase (OD_600_ = 0.8 -1.0). Subsequently, yeast cells were harvested, and total proteins were collected using a trichloroacetic acid (TCA) extraction protocol adapted from Cox et. al. ^35^ Total protein levels in yeast lysates were estimated using a BCA Protein Assay Kit (Pierce Thermofisher) according to the manufacturer’s specifications. SDS-PAGE and western blot analysis were performed using 100 µg/lane of total protein. For antigen detection, the PVDF membrane was incubated with TSU-8 (*Pd*CTR1a specific) anti-seria at a 1:2000 dilution in 1X TBST containing 1% non-fat milk powder. After incubation, the membrane was washed three times with 1X TBST for 5 minutes each. Finally, the membrane was incubated with a secondary antibody, a goat anti-rabbit secondary antibody coupled to Alexa Fluor 647 (AF647), at a 1:10,000 dilution for an hour. Excess secondary antibody was removed by washing the membrane three times with 1X TBST for 5 minutes each. Western blot images were collected using a Bio-Rad ChemiDoc MP imaging system. The monoclonal antibodies GF28R (eBioscience) and YL1/2 (Invitrogen) were used to assess total GFP and α-tubulin expression levels respectively.

Western blot detection of *Pd*CTR1a in its native *Pseudogymnoascus destructans* host was performed on *P. destructans* lysates of fungi grown on chemically defined SC-Ura growth plates.*P. destructans* spores were used to inoculate fresh SC-Ura growth plates alone or SC-Ura growth plates supplemented with 500 µM CuSO_4_ or 800 µM BCS and cultivated for 10 days at 15°C. Fungal cells were suspended in 2.0 ml of TE buffer by gently rubbing the mycelium mat with an incubation loop. After harvesting, the cells were washed twice with 1X TE buffer. Protein was extracted using the same trichloroacetic acid (TCA) protocol.^35^ A total of 110 µg of protein per lane was loaded onto an 4-12% gradient SDS-PAGE gel. Before transferring the protein to a PVDF membrane, the gels were stained with 4% trichloroethanol (TCE) solution for an hour and then washed once with deionized water.^36, 37^ The gels were imaged using the Bio-Rad stain-free gel system to assess total protein levels. Following imaging, the gels were transferred to a PVDF membrane using an iBlot 3 Western Blot Transfer System. The subsequent steps were similar to the yeast Western blotting protocol described above using TSU-8 antisera (1:2000 dilution) and a goat anti-rabbit AF647 conjugated secondary antibody.

### 2.9 ICP-MS

Yeast strains were cultured overnight in 50 mL of Sc-His media at 30°C until reaching an OD_600_ nm of approximately 0.8 to 1.0. The cells were harvested by centrifugation and washed sequentially with 1X TE buffer and then with Milli-Q water to remove residual media components and buffer salts. Subsequently, the washed cell pellet was resuspended in 1 mL of Milli-Q water, and the OD_600_ was recorded for cell density estimation. For metal ion analysis, 500 µL of the resuspended cells was mixed with 500 µL of 67-70% nitric acid (TraceMetal grade) and incubated at 75°C for 1 hour, at which time 250 µL of 30% hydrogen peroxide (H_2_O_2_) was added. The following mixture was heated to 95°C for 1 hour or until the yeast was fully digested. The volume was adjusted to 10 mL with Milli-Q water after cooling the digested solution to room temperature. The final metal concentration was determined using Agilent 8900 Inductively Coupled Plasma-Mass Spectrometer (ICP-MS), using a 23 multi-element standard IV (Centipur/ Merck KGaA, Darmstadt, Germany).

### 2.10 RT-qPCR

The fungi *P. destructans* were cultured for 10 days on 1.5% agar-supported SC-Ura media under three conditions: Control (SC-Ura), high copper (SC-Ura + 500 µM CuSO_4_), and low copper (SC-Ura + 800 µM BCS). After cultivation, cells were harvested and washed with 1X TE buffer to remove residual media components. Total RNA was extracted using the TRIZOL method according to the manufacturer’s specifications. Subsequently cDNA libraries were synthesized using the NEB LunaScript® RT SuperMix Kit following the manufacturer’s guidelines using 100 ng of total *P. destructans* RNA as an input. The resulting cDNA library was diluted 20-fold and used as a template for real-time quantitative-PCR (qPCR) analysis. A typical 20 µl qPCR reaction consisted of 10 µl of Luna Universal qPCR Master Mix (2X), 0.5 µl each of forward and reverse primers (see SI Table 1) (10 µM each, resulting in 0.25 µM final concentration each), and 2 µL of cDNA template. The qPCR cycling protocol included an initial denaturation step at 95°C for 60 seconds, followed by denaturation at 95°C for 15 seconds for 40–45 cycles. Each cycle included an extension step at 60°C for 30 seconds, with fluorescence detection using the FAM scan mode on the real-time instrument. Post-cycling, a melt curve analysis was conducted by heating from 60°C to 95°C with varied time intervals to assess PCR product specificity. The resulting data was analyzed using the ΔΔCt method using *P. destructans* actin transcripts (VC83_07844) as a reference gene standard.^38^

### 2.11 Yeast fixation for microscopy

The fixation of yeast was performed as described by Uzunova et. al.^39^ Briefly, yeast samples were grown overnight in SC-His liquid media to an OD_600_ value of 0.8 -1.0. One mL of the culture was spun at 3,000 x g at 4 °C for 10 minutes. The supernatant was discarded and the resulting yeast pellet was suspended in 100 μL of paraformaldehyde-sucrose solution (4 g/100 mL paraformaldehyde, 3.4 g/100 mL sucrose). The samples were vortexed for 5-15 seconds. The samples were then incubated at room temperature for 15 minutes. Following incubation, the samples were spun at 3,000 x g for 1 minute. The supernatant was removed, and the yeast pellet was washed using 500 μL of potassium phosphate and sorbitol solution (1.20 M sorbitol, 0.1 M potassium phosphate, pH = 7.5) and stored for up to 4 weeks at 4 °C protected from light. ProLong Gold Antifade mounting media with DAPI (Invitrogen) was added to a microscope slide. To the droplet, 2-5 μL of the stored, fixed cells were added and mixed. The cover slip was added and sealed with clear nail polish. Microscope images were collected on a Zeiss Axio Observer 7 system at 1000X total magnification equipped with an 820 monochrome digital camera, aptotome-3, and DAPI/GFP filter sets. The Zeiss Zen software package was used for image processing.

## 3. Results

### 3.1 Identification of PdCTR homologs

The family of high-affinity Copper transporters (CTRs) facilitates the trafficking of reduced Cu(I) ions across the plasma membrane using characteristic N-terminal MXXM/MXM Cu-binding motifs and three plasma membrane-spanning alpha helix domains. Analysis of the *P. destructions* genome for CTR homologs identifies two putative CTR genes, VC83_00191*(Pd*CTR1a) and VC83_04814(*Pd*CTR1b). The *Pd*CTR1a and *Pd*CTR1b isoforms are 206 and 149 amino acids in size, respectively, and display 23% sequence identity. Similar to structural similarity observed in the CTR isoforms found in *Saccharomyces cerevisiae* and *Cryptococcus neoformans, Pd*CTR isoforms display three predicted transmembrane domains and a methionine-rich region at their N-terminus (Figure 1a). There are two distinct N-terminal methionine-rich regions found in *Pd*CTR1a, which are flanking a series of flexible GSSH motifs that may assist in extracellular Cu-binding (Figure 1b).^40, 41^ Conserved methionine residues, which can assemble the Cu-methionine selectivity filter to discriminate against oxidized Cu^2+^ and reduced Cu^+^ ions in mature cell surface trimeric CTR assemblies, are present and can be found in the second transmembrane helix domain and are predicted to localize near the cell surface (Figure 1b and 1c).^41, 42^ Multiple sequence alignment of *P. destructans, S. cerevisiae*, and *C. neoformans* CTR isoforms cluster suggest *Pd*CTR1a is closely related to the high-affinity CTR1 Cu-transporter found in *C. neoformans* whereas *Pd*CTR1b is most closely related to the low-affinity Cu-transporter CTR2 found in *S. cerevisiae*(Figure 1d and Supplemental Figure 1). Together, these data support that the *P. destructans* genome encodes for two putative CTR genes that can be used to capture extracellular copper to fulfill its basic nutritional copper requirements.

**Figure 1.**
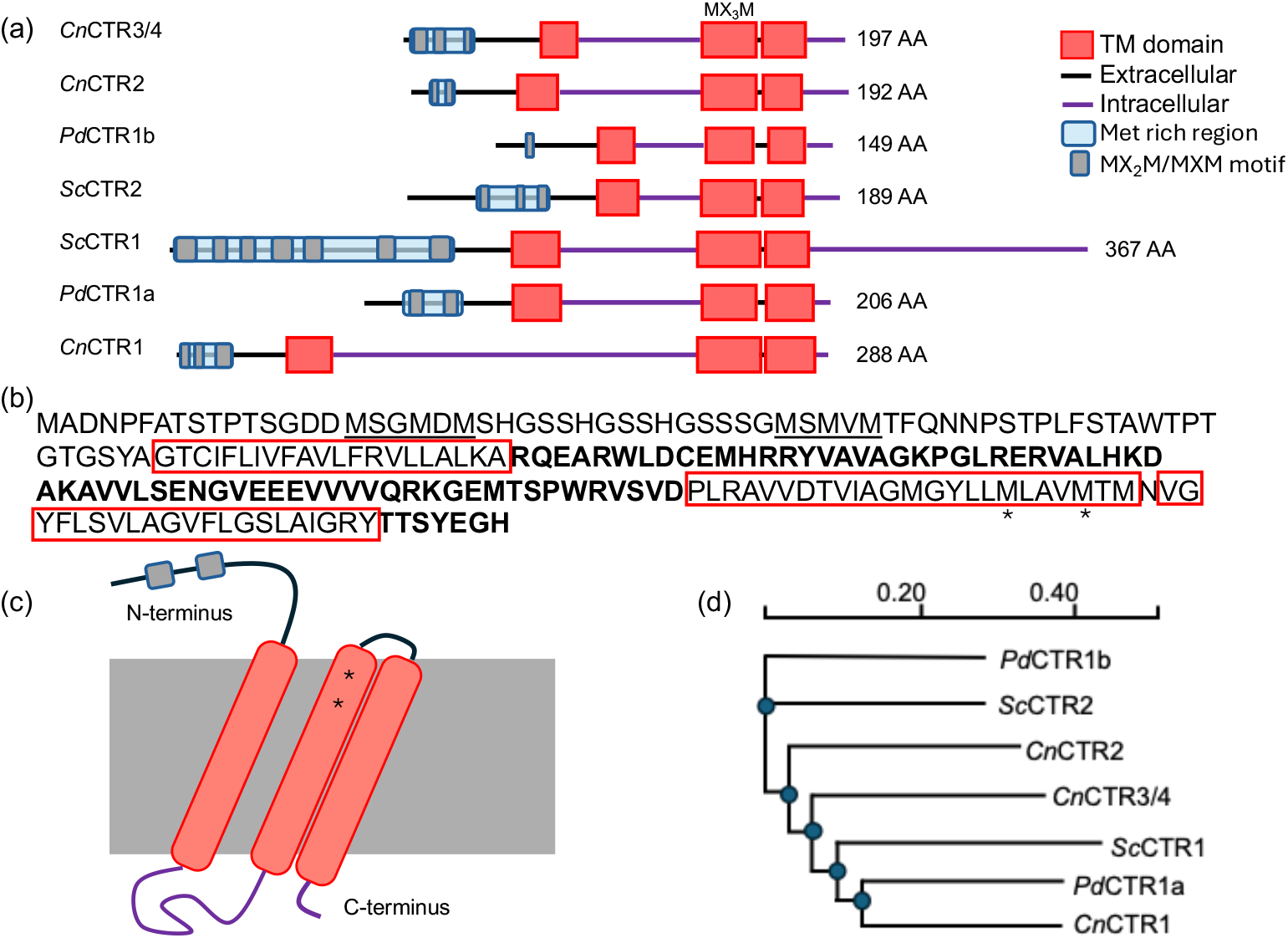
Domain architecture and similarity of *P. destructans* Cu-transporter isoforms (CTRs): (**a**) A cartoon representation of CTR isoforms found in *S. cerevisiae* (Sc), *C. neofromans* (*Cn*) and *P. destructans* (*Pd*) as predicted by DeepTMHMM. Extracellular regions are displayed as a black lines and cytosolic region as purple lines. Predicted transmembrane helixes are indicated as red boxes. Predicted extracellular Cu-binding Met-rich regions are displayed by a blue box with Met-rich domains indicated as grey ovals. (**b**) The amino acid sequence of *Pd*CTR1a. Met-rich regions are underlined and transmembrane domains are boxed in red. Cytosolic regions are indicated in bold and conserved methionine residues in the Cu-selectivity filter are indicated by a * symbol. (c) Cartoon representation of the *Pd*CTR1a isoform. (**d**) Phylogeny of *Sc, Cn*, and *Pd* CTR isoforms as predicted by Clustal omega.

### 3.2 Recombinant expression of PdCTR1a-GFP in Saccharomyces cerevisiae

To test the functionality of *Pd*CTR1 isoforms, we constructed a pRS313-based plasmid system using the native *S. cerevisiae* Ctr1 promoter and terminator to drive the recombinant expression of cytosolic GFP or the *Pd*CTR1a-GFP fusion protein (see Supplemental file 2 for plasmid information). This strategy relies on the Cu-sensing MAC1 transcription factor to drive recombinant expression only under Cu-deficient growth conditions, thereby, avoiding the transcription of recombinant proteins and when cellular Cu-quotas are sufficient.^43^ Flow cytometry analysis of *S. cerevisiae* BY4741 (WT) and BY4741*ctr1Δ* (*ctr1*Δ) yeast transformed with empty vector (i.e. pRS313) or the recombinant expression of cytosolic GFP or *Pd*CTR1a-GFP driven by the Ctr1 promoter display distinct fluorescence profiles when cultured under different copper stress conditions in Synthetic Complete-Histidine (SC-His) growth media (Figure 2). In all experimental conditions, logarithmic yeast cultures harboring GFP expression cassettes exhibit an approximate 5 – 100-fold increase in GFP fluorescence intensity when compared to the empty vector controls. The highest levels of GFP fluorescence were found in yeast grown in the presence of 100 μM of the copper chelator BCS (Figure 2c). Both WT and *ctr1Δ* yeast backgrounds display comparable expression levels for each plasmid construct, with cytosolic GFP expression strains displaying the highest GFP fluorescence levels. This suggests that the Cu-sensing MAC1 transcription factor is active and the Ctr1 promoter can drive recombinant GFP construct across the two yeast backgrounds. However, supplementing SC-His growth media with 50 μM copper sulfate leads to minimal GFP fluorescence profiles in both WT and *ctr1Δ* yeast when compared to empty vector control strains (Figure 2b). This implies that cellular copper levels are sufficient to repress MAC1 transcription factor binding to the Ctr1 promoter under growth conditions supplemented with excess copper sulfate in both the WT and *ctr1Δ* yeast backgrounds. Further growth experiments in standard SC-His growth media alone reveal that only *ctr1Δ* harboring the cytosolic GFP construct display elevated levels of GFP fluorescence (Figure 2c). Whereas WT yeast harboring both cytosolic GFP and *Pd*CTR1a-GFP constructs, as well as BY4741ctr1Δ harboring the *Pd*CTR1a-GFP construct, display lower GFP fluorescence levels. The ability of *Pd*CTR1a-GFP within a *ctr1Δ* yeast background to yield similar levels of GFP found in WT yeast strains suggests *Pd*CTR1a-GFP can restore intracellular Cu levels closer to wild-type levels and is a functional copper permease.

**Figure 2.**
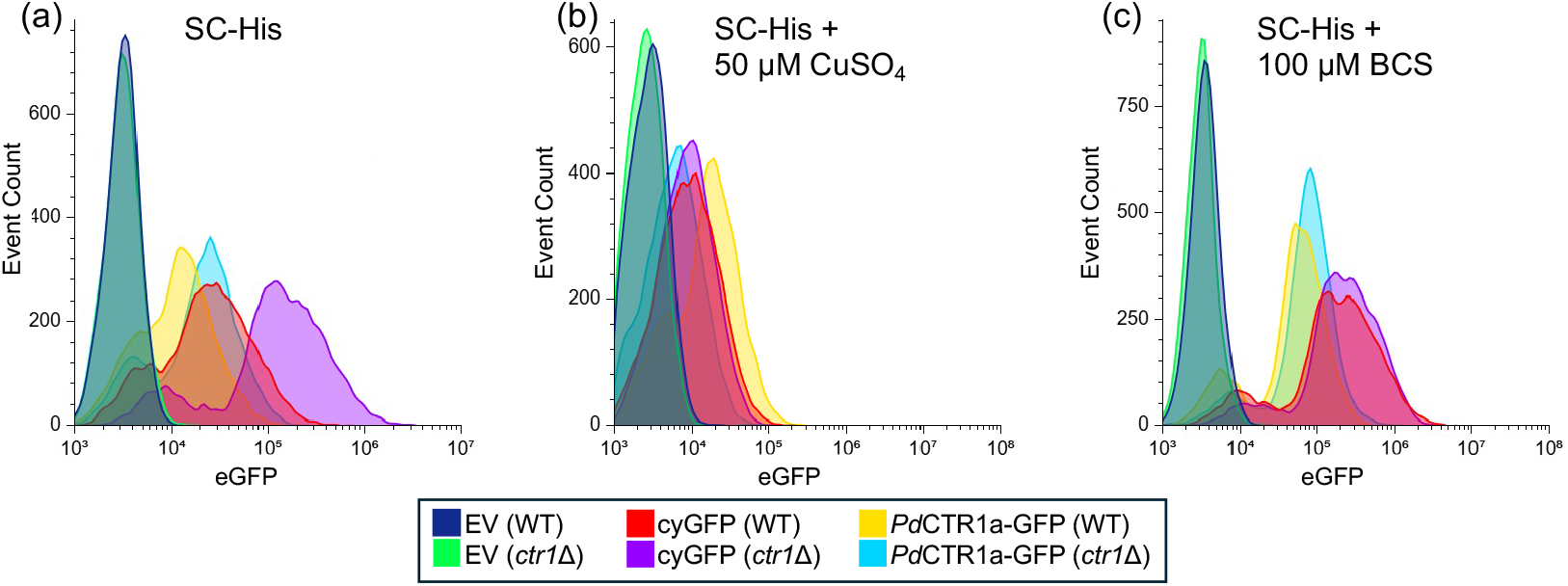
Flow cytometry of recombinant *Saccharomyces cerevisiae* strains. GFP fluorescence profiles of BY4741 (WT) and BY4741*ctr1*Δ (*ctr1*Δ) yeast harboring empty vector (EV), cytosolic-GFP (CyGFP) or *Pd*CTR1A-GFP under the control of the *S. cerevisiae* Ctr1 promoter and terminator. Yeast was grown to the log phase in SC-His under varying Cu-stress conditions: (a) control (i.e., SC-His), (**b**) SC-His + 50 μM Cu-sulfate, (**c**) SC-His + 100 μM BCS. A total of 10,000 events are displayed per trace. Traces are normalized to the background fluorescence of yeast harboring the pRS313 empty vector plasmid alone set to an intensity of 1 × 10^3^ eGFP fluorescent units.

The localization of patterns of *Pd*CTR1a-GFP fusion protein was determined using fluorescence microscopy on fixed WT and *ctr1Δ* yeast samples grown in SC-His supplemented with 100 μM BCS (Figure 3). Imaging of the empty vector or plasmids encoding for the recombinant expression of cytosolic GFP control strains was also performed (Figure 3). Under these Cu-limiting growth conditions, we envision the localization of *Pd*CTR1a-GFP could reside in two primary locations: 1. the cell surface plasma membrane or 2. An intracellular vacuole or cellular compartment. In *Pd*CTR1a-GFP expressing strains, we observe diffuse and low levels of GFP fluorescence that localize to the perinuclear region and cellular surface compartment. These *Pd*CTR1a-GFP localization patterns are distinct from the cytosolic GFP expression strains that display large levels of GFP fluorescence signals and are located throughout the cytoplasm. Together, this data indicates that *Pd*CTR1a-GFP fusion protein is stably expressed in BY4741 yeast and the *Pd*CTR1a protein is associated with plasma membranes.

**Figure 3.**
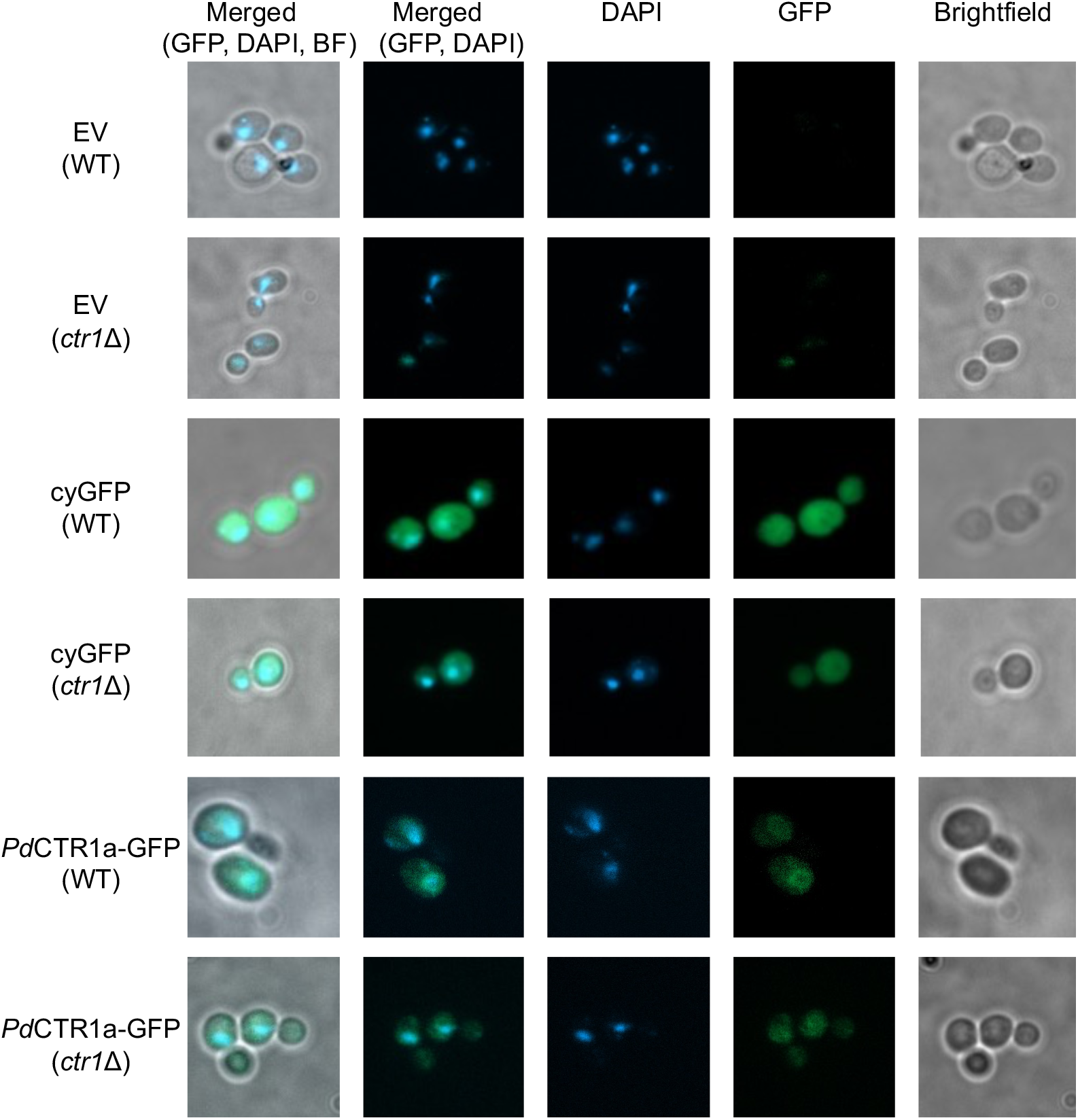
Fluorescent microscopy images of recombinant *Saccharomyces cerevisiae* strains. Images of fixed yeast samples taken under brightfield conditions as well as single channel fluorescent images using GFP and DAPI filter sets were performed at 1000X total magnification. Single and merged multi-channel images are displayed as indicated on the top legend. The left legend indicates the recombinant expression plasmid, and the yeast background is indicated below in parenthesis.

To validate the expression fidelity of the *Pd*CTR1a-GFP fusion protein, we examined the cellular contents of all yeast strains constructed under Cu-limited growth conditions by western blot. Based on flow cytometry analysis, we anticipated observing similar levels of cytosolic GFP or *Pd*CTR1a-GFP protein across the WT and *ctr1Δ* yeast backgrounds. In fact, we observe a cytosolic GFP protein band migrating at approximately 34 kDa and a band corresponding to the *Pd*CTR1a-GFP fusion protein migrating at approximately 55 kDa when probed with an anti-GFP specific antibody (Figure 4b). The successful expression of the *Pd*CTR1a-GFP fusion protein is further supported by western immunoblot detection of the *Pd*CTR1a-GFP fusion protein using anti-*Pd*CTR1a serum against raised against *Pd*CTR1a residues K86-Y102 and G107-K122 (Figure 4b). These data, in conjunction with the flow cytometry and imagining studies, support the notion that *S. cerevisiae* can serve as a recombinant host for the study of *Pd* copper transporters.

**Figure 4.**
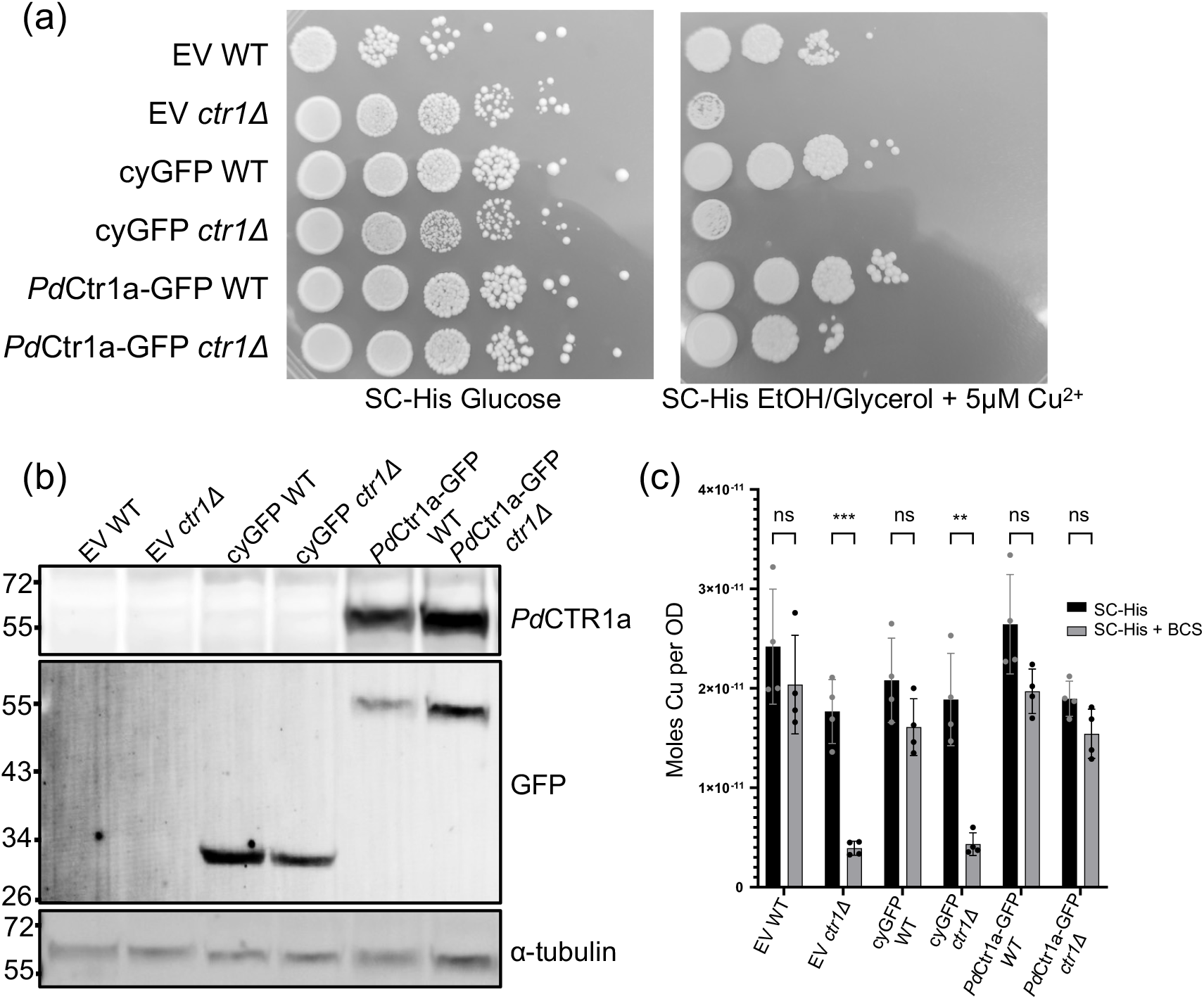
Characterization of *S. cerevisiae Pd*CTR1a-GFP expressing strains: (**a**) *Pd*CTR1a-GFP partially rescues the Cu-respiratory growth defect of *ctr1*Δ yeast. Wild-type (WT) and *ctr1*Δ strains were grown on SC-His selection media using glucose (Left) or glycerol/ethanol (Right) as a sole carbon source. Strains harboring the empty vector (EV), cytosolic GFP (CyGFP), and *Pd*CTR1a-GFP fusion are shown. Ten-fold serial dilutions of yeast are displayed and imaged after 4 days (Glucose) and 9 days (Glycerol/Ethanol). (**b**) Western Immunoblot of WT and *ctr1*Δ containing strains harboring the EV, cyGFP, and *Pd*CTR1a-GFP expression plasmids under SC-His + BCS growth conditions. Bottom panel probes for α-tubulin as a loading control. Middle panel probes for GFP and exhibits signals for both cyGFP and *Pd*CTR1a-GFP expression strains. Top panel probes for *Pd*CTR1a using TSU8 anti-serum. Protein molecular weight markers (kDa) are displayed on the left of each blot. (**c**) Yeast total cellular copper levels as determined by inductively coupled plasma mass spectrometry (ICP-MS). (**d**) Significance was determined by ANOVA with multiple unpaired t-tests. Differences were considered significant at *p* < 0.05. Significance level was ** = *p* < 0.01 and *** = *p* < 0.001.

### 3.3 PdCTR1a-GFP complements Saccharomyces cerevisiae BY4741ctr1Δ to increase Cu-fitness under Cu-stress conditions

Due to the similarities between *Pd*CTR1a isoforms and other high-affinity CTR homologs, we tested the ability of *Pd*CTR1a to complement the Cu-respiratory growth defect in *ctr1Δ* yeast (Figure 4a).^40, 41, 44, 45^ Whereas both WT and *ctr1Δ* yeast grow efficiently on glucose media, when grown on nonfermentable carbon sources (i.e., EtOH/glycerol), *ctr1Δ* yeast cannot fulfill the cellular copper quota needed to efficiently load cytochrome C oxidase for efficient respiration. ^*46*^ Poor growth behavior is even observed in BY4741*ctr1Δ* yeast, with an additional 5 μM copper sulfate added to SC-His EtOH/glycerol growth conditions. However, the expression of the *Pd*CTR1a-GFP in *ctr1Δ* yeast can partially rescue this Cu-respiratory growth defect (Figure 4a), presumably by increasing total cellular copper levels. To test this notion, we quantified the total cellular copper levels of all yeast strains constructed under Cu-replete and Cu-limited growth conditions. No significant difference in cellular Cu levels are found between WT and *ctr1Δ* yeast when they are grown under Cu-replete growth conditions. All strains display an average cellular copper level of approximately 2 × 10^−11^ moles Cu/OD (Figure 4c). However, when challenged with low levels of the extracellular copper chelator BCS, *ctr1*Δ yeast harboring empty vector and cytosolic GFP vector exhibit approximately a 4-fold reduction in cellular copper levels. This behavior is not observed in the *ctr1Δ* yeast that is capable of expressing *Pd*CTR1a-GFP where total cellular copper content remains near wild-type levels suggesting that the *Pd*CTR1a-GFP is functionally redundant to *S. cerevisiae* Ctr1p under the tested Cu-stress growth conditions.

### 3.4 Characterization of PdCTR1 isoforms in P. destructans under Cu-stress growth conditions

The presence of two CTR1 isoforms, *Pd*CTR1a and *Pd*CTR1b, in the *P. destructans* genome, suggests they have the capacity to participate in fungal copper homeostasis, but their roles in their native fungal host are not defined. Thus, we next turned to characterize the transcriptional response of *Pd*CTR1a and *Pd*CTR1b genes in *P. destructans* under copper stress growth conditions using chemically defined SC-Ura growth media. The transcriptional regulation of *Pd*CTR1a and *Pd*CTR1b genes under control, high copper, and Cu-withholding growth conditions using *Pd*Actin (VC83_07844) as an internal reference standard is displayed in Figure 5a. Here both *Pd*CTR1 isoforms display similar trends in transcriptional control based on copper bioavailability. We find that adding the copper chelator BCS leads to a 7.1 increase in *Pd*CTR1a transcripts and a 6.5 increase in *Pd*CTR1b transcripts levels versus control conditions. Whereas, the addition of copper sulfate to the growth media represses both *Pd*CTR1a and *Pd*CTR1b total transcript levels. The *Pd*CTR1a isoform, however, is more severely impacted by added copper. An approximate 33-fold reduction in *Pd*CTR1a transcripts and 2.5-fold decrease reduction in *Pd*CTR1b transcripts, respectively, are found versus control SC-Ura growth conditions. Together, this data suggests that the *Pd*CTR1a and *Pd*CTR1b transcripts are regulated by a copper-sensing transcription factor.

**Figure 5.**
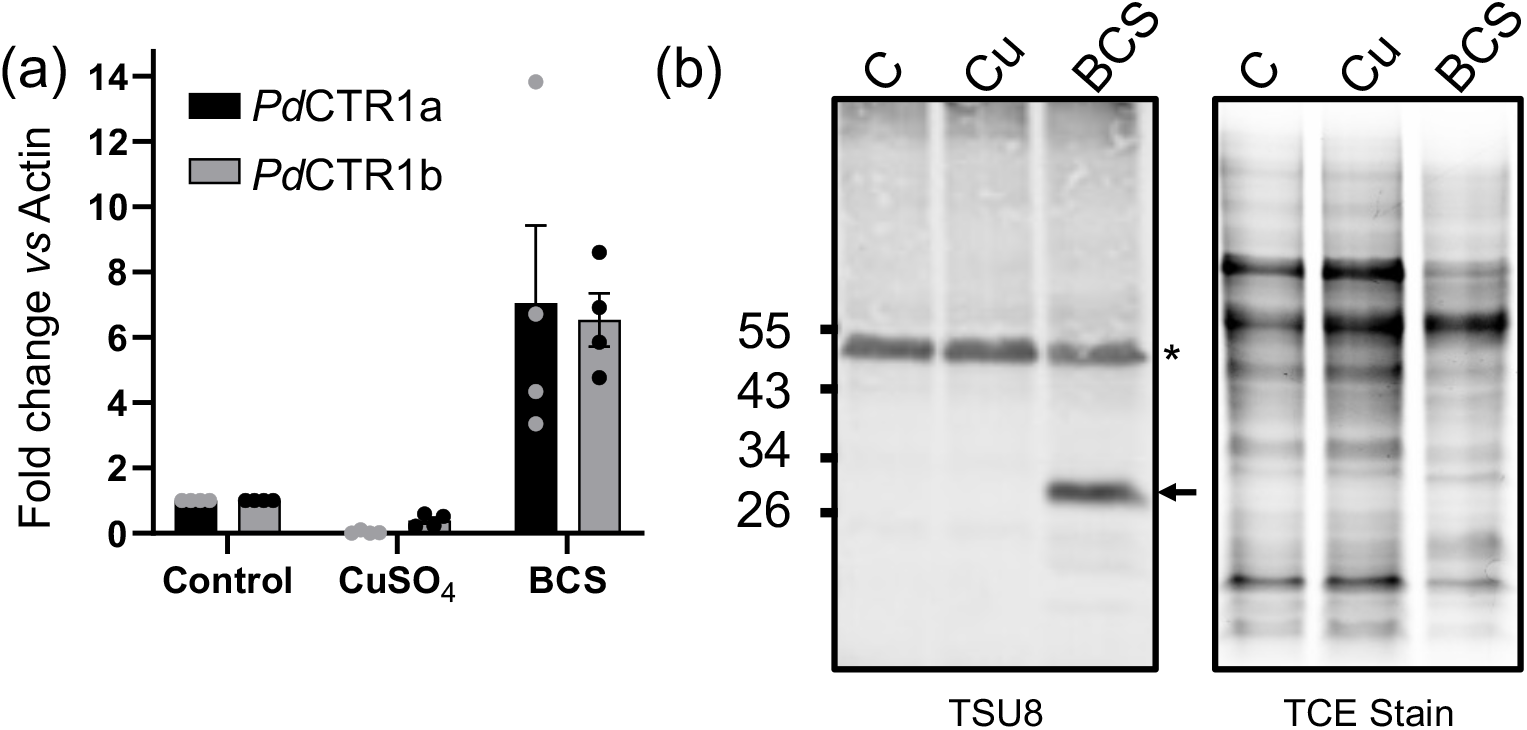
The CTR1 isoform *Pd*CTR1a are transcriptionally regulated by extracellular bioavailable Cu levels and expressed in Cu-starved *Pd* cells. (**a**) Real-time quantitative PCR (RT-qPCR) of *Pd*CTR1a (black bars) and *Pd*CTR1b (grey bars) isoforms under Cu-stress growth conditions (see experimental section). Transcript levels are reported using the ΔΔCT method using *Pd*Actin (VC83_07844) as the internal housekeeping gene. Experiments represent four biological replicates with the control levels normalized to 1.0.; (**b**) Isoform *Pd*CTR1a protein is present in Cu-starved *Pd* cells. The left panel is a western immunoblot of *Pd* lysates using TSU8 anti-*Pd*CTR1a antiserum (see experimental section). * represents a non-target peptide found in all *Pd* samples in the TSU8 antiserum and the assigned *Pd*CTR1a is indicated by an arrow. The right panel is the TCE stained gel image used to quantity equivalent protein loading levels of *Pd* lysates. Lane legends “C” -control (i.e. SC-Ura), “Cu”-SC-Ura + 500 μM Cu-sulfate, “BCS” -SC-Ura + 800 μM BCS.

Based on the recombinant *Pd*CTR1a-GFP expression experiments performed in *S. cerevisiae*, there was the opportunity to use the validated *Pd*CTR1a anti-serum to probe *Pd*CTR1a expression levels in its native *P. destructans* fungal host. We performed western immunoblot analysis of *P. destructans* whole cell lysates using TSU8 *Pd*CTR1a anti-serum to probe for *Pd*CTR1a expression levels and used TCE staining to assess total protein loading levels (Figure 5b). Probing with TSU8 serum, we identify a consistent band at ∼ 55 kDa that is found in all *P. destructans* lysates. However, when *P. destructans* is cultured with the Cu-chelator BCS, we identified a new and unique protein band at ∼ 30 kDa, which we attribute to *Pd*CTR1a protein. Taken together with the qRT-PCR experiments in *Pd, Pd*CTR1a is a biomarker for Cu-withholding stress at both the transcriptional and translational levels.

## 4. Discussion

The family Cu-transporters (CTRs) play important roles in the import of essential Cu-micronutrients across lipid bilayers into the cytosolic of eukaryotic cells from yeast to man.^47, 48^ To adapt and survive in animal hosts, many infectious fungal pathogens often have redundant high-affinity CTR transporters to scavenge copper from extracellular copper pools to fulfill their basic copper metabolic requirements over the course of host colonization and infection.^14^ Thus, CTR transporters, along with several copper metalloenzymes, including Cu/Zn-SODs^49-51^ and laccases^52, 53^ which the copper CTR transporters supply, are important virulence factors in many animal infecting fungal pathogens. Recent RNA-Seq studies on active *P. destructans* infection sites of white-nose syndrome bats suggest *P. destructans* is starved of trace metal micronutrients.^21, 22^ The presence of elevated transcripts of S100 proteins suggests the bat host may be employing a mechanism of nutritional immunity to limit essential metal micronutrient availability to limit *P. destructans* growth. Similar transcriptomic studies aimed at describing how the *P. destructans* transcriptome changes during bat-host colonization have identified *Pd*CTR1a as an essential virulence factor to assist in trace copper acquisition.^22^ This paper describes the biochemical characterization of the virulence factor *Pd*CTR1a using *S. cerevisiae* as a model system and in the *P. destructans* native host.

Baker’s yeast *Saccharomyces cerevisiae* has proven to be an excellent model fungus to study basic metal cell biology. The genome of the laboratory BY4741 yeast strain encodes for two functional CTR transporters: Ctr1p and Ctr2p. The Ctr1p transporter is the primary high-affinity Cu transporter that supplies most copper import during low copper growth conditions and contains a C-terminal regulon^54^ that is degraded in the presence of high copper levels.^55^ Whereas Ctr2p primarily localizes to the vacuole^56^ and may assist in mobilizing intracellular copper pools or as a low-affinity Cu-transporter when localized to the plasma membrane.^57^ Localization studies with recombinant *Pd*CTR1a-GFP suggest that *Pd*CTR1a is trafficked as a functional Cu-transporter to the *S. cerevisiae* cell surface plasma membrane to deliver copper to the cytoplasm. The ability to rescue the Cu-respiratory growth defect in *ctr1Δ* yeast corroborates this cell surface localization pattern and our ICP-MS studies with *ctr1Δ* yeast grown in the presence of the Cu-chelator BCS validates *Pd*CTR1a copper trafficking activity. Our observations describing *PdCTR1a-GFP* activity in BY4741*ctr1Δ* yeast are comparable to other studies that have been used to assign a function to high-affinity Cu-transporters from other eukaryote species. ^40, 41, 44, 45, 58^ However, our experiments *Pd*CTR1a-GFP require low levels of supplemental Cu (i.e. 5 μM) to restore this respiratory growth defect. We interpret these results as our recombinant expression construct using the *S. cerevisiae* Ctr1 promoter results in lower levels of recombinant protein to the strong constitutively GPD promoter used in previous studies.^40, 42-44^ An alternative explanation for the need for low levels of supplemental copper could be that *Pd*CTR1a has a lower affinity for Cu than the native *S. cerevisiae* Ctr1p or that an additional protein cofactor is needed to assist in Cu-delivery to *Pd*CTR1a. In *Cryptococcus neoformans*, the glycophosphatidyl inositol (GPI) anchor protein Bim1p was shown to assist in high-affinity Cu delivery to its Ctr1 homolog.^59^ Surprisingly, the *P. destructans* genome also encodes for three Bim1p homologs, and one or more Bim1P homologs may be needed to assist in boosting *Pd*CTR1a Cu-trafficking activity.

Our study also describes the transcriptional regulation of *Pd*CTR1a and *Pd*CTR1b isoforms in the native *P. destructans* fungus. We find that both CTR isoforms are transcriptionally active and likely participate in the Cu-stress response. In *Saccharomyces cerevisiae*, the regulation of ctr1 and ctr2 transcripts are controlled through different transcription factors. High affinity CTR1 transcript levels are regulated through the Cu-sensing Mac1^43^ transcription factor under copper deficiency, whereas CTR2 transcript levels respond to iron and copper withholding stress.^60, 61^ Our data is consistent with the notion that both *Pd*CTR1a and *Pd*CTR1b isoforms respond to copper withholding stress in the native *P. destructans* fungus. However, the identity of this transcription factor is not clear. We cannot identify Cu-binding consensus sequences for MAC1 (5’-TTTGC(T/G)C(A/G)-3’)^62^ and CUF1 (‘5-AA(T/G)GGC(T/G)C-3’)^63^ nor CuSE (5’-D(T/A)DDHGCTGD-3’)^64^ in the *Pd*CTR1a and *Pd*CTR1b promoter region. This may infer that *P. destructans* uses another Cu-sensing transcription factor(s) or an alternative Cu-responsive element sequence. Further studies are needed to identify the transcription factors or elements that regulate *Pd*CTR1a and *Pd*CTR1b transcription levels.

In the native *P. destructans* host, we were also able to detect *Pd*CTR1a protein under conditions that correlated with the transcriptional regulation patterns observed in our RT-qPCR study. We observe a unique band at approximately 30 kDa, which we assign to *Pd*CTR1a when *P. destructans* is cultured with the Cu-chelator BCS. While we anticipate the primary driver for high-affinity CTR transporters is at the transcriptional level, regulation of CTR activity and stability can be influenced at the post-translational level, specifically at the C-terminus.^54, 65^ Previous studies with *S. cerevisiae* showed that CTR1p turnover is largely regulated by the E3 ubiquitin ligase Rsp5 targeting two lysine residues at the c-terminal region in response to sudden cellular copper flux.^55^ While the predicted c-terminal domain of *Pd*CTR1a does not encode for any lysine residues past the third predicted transmembrane, there are several potential lysine residues in the cytosolic region between the first and second transmembrane that may participate in modulating *Pd*CTR1a protein turnover in the native *P. destructans* fungus. Additionally, both *Pd*CTR1a and *Pd*CTR1b isoforms contain the possible copper metal binding sequences, EGH and ACH, respectively at their C-terminus, which may facilitate Cu-delivery to Cu-chaperones such as Atox1.^65^ Further studies into the mechanisms of *Pd*CTR1a protein trafficking and degradation are important for *P. destructans* copper homeostasis and they should be addressed in future studies.

## 5. Conclusions

In summary, this report describes the biochemical and metal cell biology of *P. destructans Pd*CTR1a which has been implemented as an essential fungal virulence factor. Recombinant studies in *S. cerevisiae* confirm that *Pd*CTR1a is a bonafide high-affinity copper transporter that can localize to the cell surface plasma to import essential Cu-micronutrients. Complementary studies in the native *P. destructans* fungus provide evidence that *Pd*CTR1a transcripts are regulated by copper bioavailability in the growth media and *Pd*CTR1a protein can be detected in *P. destructans* cultured under Cu-restrictive growth conditions. Our data supports the notion that *Pd*CTR1a can be used as a biomarker for *P. destructans* Cu-stress in White Nose Syndrome samples.

## Author Contributions

Conceptualization, A.D.F., S.A. and R.L.P; Methodology, A.D.F., S.A. and R.L.P; Validation, A.D.F., S.A. and R.L.P; Formal analysis, A.D.F., S.A. and R.L.P; Investigation, A.D.F., S.A. and R.L.P and C.E.M.; Resources, A.D.F., S.A. and R.L.P.; Data curation, A.D.F., S.A. and R.L.P; Writing—original draft, A.D.F., S.A. and R.L.P; Writing—review & editing, A.D.F., S.A. and R.L.P; Visualization, A.D.F., S.A. and R.L.P. All authors have read and agreed to the published version of the manuscript.

## Data Availability Statement

The data presented in this study are available from the corresponding author upon reasonable request.

Acknowledgments

We would also like to thank Dr. L.K. Lewis for the yeast strains and plasmids used in this study. A.D.F., S.A. and R.L.P. would like to thank the Texas State University for financial support.

## Supplemental File 1

Multiple sequence alignments of CTR homologs from *P. destructans, S. cerevisiae, and C. neoformans*.

## Supplemental File 2

Information on the plasmids used in this study.

## Supplemental Table

List of primers used in this study.

