## Supplemental File 1 for "Characterization of a High-Affinity Copper Transporter in the White-Nose Syndrome Causing Fungal Pathogen *Pseudogymnoascus destructans*"

**Supplemental File S1.** BLAST sequence alignment of Cu-transport homologs from *S. cerevisiae (Sc)*, *C. neoformans (Cn)*, and *P. destructans (Pd).* Protein sequences were retrieved using the identifiers YPR124W (*Sc*CTR1), YHR175W(*Sc*CTR2), CNAG_00979(*Cn*CTR2), CNAG_01872 (*Cn*CTR2), CNAG_07701(*Cn*CTR1), VC83_00191 (*Pd*CTR1a), and VC83_04814(*Pd*CTR1b). Large portions of non-homologous alignment sequence are designated by a bold "X" with the subscript number corresponding to the number of un-aligned amino acids.

```
ScCTR1                  MEGMNMGSSMNMDAMSSASKTVASSMASMSMDAMSSASKTILSSMSSMSMEAMSSASKTL    60
PdCTR1a                 ------------------------------------------------------MADNPF    6
CnCTR1                  ------------------------------------------------------------    0
CnCTR2                  ------------------------------------------------------------    0
CnCTR3/4                ------------------------------------------------------------    0
PdCTR1b                 ------------------------------------------------------------    0
ScCTR2                  ------------------------------MDDKKT-----------WSTVTLRTFNQLV    19

ScCTR1                  ASTMSSMASMSMGSSSMSGMSMSMSSTPTSSASAQTTSDSSMSGMSGMSSSDNSSSSGMD    120
PdCTR1a                 A------TSTPTSGDDMSGMDM-------------------SHGSSHG---SS-----HG    33
CnCTR1                  --------------MDMSGMD----------------------GMDHM---SS-----TS    16
CnCTR2                  -----------------------MNHGD-----------HSKHTMPDM---D--------    15
CnCTR3/4                --------------MDMGNMGMGMGMGM-----------DSGHNHSHM---NMGSGHGAD    32
PdCTR1b                 -------------------------MSH-----------SMGHG-------DHD-----A    12
ScCTR2                  T------SSLIGYSKKMDS--MNHKMEG-----------NAGHDHSDM---HMG-----D    52

ScCTR1                  MDMSMGMNYYLTPTYKNYPVLFHHLHANNSGKAFGIFLLFVVAAFVYKLLLFVSWCLEVH    180
PdCTR1a                 SSSGMSMVMTFQNNP-STPLFSTAWTPTGTGSYAGTCIFLIVFAVLFRVLLALKARQEAR    92
CnCTR1                  SNMSMSMKMYFHGTIGGDLLWFASWMPSSAGATVGVCIGLFILAIFERYLVAFRRACDAA    76
CnCTR2                  -MPACSMNMLWNNQVADTCVVFRSWHISGTWTMILSCLIIIGISVFYSYLLHYIKDYDRH    74
CnCTR3/4                SGHACRISMLLNFNTVDACFLSPNWHIRSKGMFAGSIIGIFFLCVLIELIRRLGREFDRW    92
PdCTR1b                 SAARCNMNMLFTWSTQDLCIIFRSWHITGPITLTISLLAIVALVAGFEALRATTARYDAA    72
ScCTR2                  GDDTCSMNMLFSWSYKNTCVVFEWWHIKTLPGLILSCLAIFGLAYLYEYLKYCVH-----    107
                              :         .  .                 : :.        :

ScCTR1                  WFKKWDKQNKY-------------STLPSANS-------KDEGKHY--------DTE---    209
PdCTR1a                 WLDCEMHRRYVAV-------AGK------PGLRERVALHKDAKX_4---------------    122
CnCTR1                  WRRGQVGYVRPCSNGPLVFSSGKSTSLPPPVLFNRRSSTKKEKDVYNPLTPSDYALEX_63-    136
CnCTR2                  VAAA----IYSST---------------QQGRRDRDGSPA----ETGLIPPIPTAYG---    108
CnCTR3/4                ----AGVNS---------------TCGELS---------------SVAEYGKD---  115
PdCTR1b                 LLKR----RDELP---------------HEELAE---------------TT---------    89
ScCTR2                  -----------------------------KRQLS---------------QR---------    114

ScCTR1                  --NNFEIQG-LPKLPNLLSDIFVP-------SLMDLFHDIIRAFLVFTSTMIIYMLMLAT    259
PdCTR1a                 SENGVE------------EEVVVVQRKGEMTSPWRVSVDPLRAVVDTVIAGMGYLLMLAV    174
CnCTR1                  KEKAVERGLVHSHLPKAVRRSLDPGREGRWSRPFRLAVDVPRGLLQALQTLIHYLLMLVV    256
CnCTR2                  GIEVGAL-------------NRI-GVTRLPLR--LRLIRAGLYAVTVAISFWLMLVA  149
CnCTR3/4                ---GAQGGAV-------------VRVAPRYVPSWP--HQILRGFIYGSQFTAAFFVMLLG    157
PdCTR1b                 --------TL-------------LPGQQQSLRDVR--AKVVRSALYGLETFYAFMIMLLF    126
ScCTR2                  --------VL-------------LPNRSLTKINQA--DKVSNSILYGLQVGFSFMLMLVF    151
                                                                 .. :         : :**

ScCTR1                  MSFVLTYVFAVITGLALSEVFFNRCKIAMLKRWDIQREIQKAKSCPGFX_98----------    319
PdCTR1a                 MTMNVGYFLSVLAGVFLGSLAIGRYTTSY-----------EGH-----------------    206
CnCTR1                  MTFNIWWMISVVIGCGVGEMLFGRFGSSH-----------VGH-----------------    288
CnCTR2                  MTYNTYLFSSIVVGAFFGHVIYEDEMDVGAVLSGTS---GKGLACH--------------    192
CnCTR3/4                MYFNVIVLIFIFLGQTVGYMLFGRDTCGGGFDFG-----AQGRCC---------------    197
PdCTR1b                 MTYNGQVMIAVGIGAFVGHLAFGGATTA-----------TRETACH--------------    161
ScCTR2                  MTYNGWLMLAVVCGAIWGNYSWCTSYSPE---ID-----DSSLACH*-------------    189
                        *      .  :  *   .
```
