## Supplemental File 2 for "Characterization of a High-Affinity Copper Transporter in the White-Nose Syndrome Causing Fungal Pathogen *Pseudogymnoascus destructans*"

Supporting Information File 2

Plasmid pAF23 (pCTR1-yeGFP) tCTR1 (addgene plasmid #223171)

*Legend*

CTR1 Promoter UNDERLINED

GFP

CTR1 Terminator **BOLD**

*Plasmid Sequence:*

catgtgtatatatgtatacctatgaatgtcagtaagtatgtatacgaacagtatgatactgaagatgacaaggtaatgcatcattctatacg
tgtcattctgaacgaggcgcgctttcctttttctttttgctttttctttttttctcttgaactcgacggatctatgcggtgtgaaataccgcacaga
tgcgtaaggagaaaataccgcatcaggaaattgtaagcgttaatattttgttaaaattcgcgttaaatttttgttaaatcagctcattttttaac
caataggccgaaatcggcaaaatcccttataaatcaaaagaatagaccgagatagggttgagtgttgttccagtttggaacaagagtc
cactattaaagaacgtggactccaacgtcaaagggcgaaaaaccgtctatcagggcgatggcccactacgtgaaccatcaccctaa
tcaagttttttggggtcgaggtgccgtaaagcactaaatcggaaccctaaagggagcccccgatttagagcttgacggggaaagccg
gcgaacgtggcgagaaaggaagggaagaaagcgaaaggagcgggcgctagggcgctggcaagtgtagcggtcacgctgcgc
gtaaccaccacacccgccgcgcttaatgcgccgctacagggcgcgtccattcgccattcaggctgcgcaactgttgggaagggcgat
cggtgcgggcctcttcgctattacgccagctggcgaaagggggatgtgctgcaaggcgattaagttgggtaacgccagggttttccca
gtcacgacgttgtaaaacgacggccagtgaattgtaatacgactcactatagggcgaattggagctccaccgcggtggcggccgctc
tagaactagtcacagacattaacccacagtacagacactgcgacaacgtggcaattcgtcgcaatacaacgacacaagtccgattg
ttcctcttcaggagcttcctgaaccaaactttttccgcaaggccgcattttgaaccgtattttgctcgttccagcctttccacgtttttgttatcta
agcaacttggcacatttccctactatactacaaaccgatacgtaaatacttccctaaatagcatatgaattattcagtaatttttaaggatcg
aaactgcacctcaactattcgttactgtggttatgttctcatgtattgatgcaaatcatgggatatttgctcaagacgacggtaaaatgagc
aaaaatggcacgatcctgaaaagagcacttttcaagattcgggctacaaaatgcaacataaaaaatgttgtattgtcatctcgagagg
gtcttgtatgttttattcctcttatgattagttcacattagtaaaacagatacgcagtgtgctcttaataaacaactactccatagctttatttgca
taacaaaacttttaagcacaaacttaaacaggtggagtaatagttcggcggcgactcaaattacatttgttggaagaatcgaatagaa
aataaaaaaaagtgtattatatttgacattcaaatatgaaaggtgaagaattattcactggtgttgtcccaattttggttgaattagatggtg
atgttaatggtcacaaattttctgtctccggtgaaggtgaaggtgatgctacttacggtaaattgaccttaaaatttatttgtactactggtaa
attgccagttccatggccaaccttagtcactactttcggttatggtgttcaatgttttgcgagatacccagatcacatgaaacaacatgactt
tttcaagtctgccatgccagaaggttatgttcaagaaagaactatttttttcaaagatgacggtaactacaagaccagagctgaagtcaa
gtttgaaggtgataccttagttaatagaatcgaattaaaaggtattgattttaaagaagatggtaacattttaggtcacaaattggaataca
actataactctcacaatgtttacatcatggctgacaaacaaaagaatggtatcaaagttaacttcaaaattagacacaacattgaagat
ggttctgttcaattagctgaccattatcaacaaaatactccaattggtgatggtccagtcttgttaccagacaaccattacttatccactcaa
tctgccttatccaaagatccaaacgaaaagacagaccacatggtcttgttagaatttgttactgctgctggtattacccatggtatggatg
aattgtacaaaggatc**ctaataactagacaaggttcgtccacctatatcttcttcttccaatatctctatacatcaaaataagaata**
**tcgttcatttctcagaagctcaaaaaaaatctcaaaatggatacttctaatgccctcatgaagaggaaagaaaataatagaca**
**cagcgaccttcctataaataacagacttgtatatagttgaatgtctaagtaattttaattcaaaaatccttaactatattaccacctt**
**gcaataataagagtgaagtgttaaaaatgatacttgcttgccttcatcgagttaccgttgtctcccacgtttctttttctcgttgga**
**cagaattgcttccgcgcgaacattttccacttttatttgaaagaggacattattcagatcgtgtcagtctcatctaattggcaaag**
**tggcaattttcgaaataacaagataactgcataaagtaacacttgtcgaattgaaagtattttgccagtgatatttaggttcgagt**
**aaagaaaatttcataaagaaatcaacaagacacaatgctg**gaaatctgctcgtcagtggtgctcagaattcgatatcaagcttatc
gataccgtcgacctcgagggggggcccggtaccagcttttgttccctttagtgagggttaatttcgagcttggcgtaatcatggtcatagct
gtttcctgtgtgaaattgttatccgctcacaattccacacaacatacgagccggaagcataaagtgtaaagcctggggtgcctaatgagt
gagctaactcacattaattgcgttgcgctcactgcccgctttccagtcgggaaacctgtcgtgccagctgcattaatgaatcggccaacg
cgcggggagaggcggtttgcgtattgggcgctcttccgcttcctcgctcactgactcgctgcgctcggtcgttcggctgcggcgagcggt
atcagctcactcaaaggcggtaatacggttatccacagaatcaggggataacgcaggaaagaacatgtgagcaaaaggccagca
aaaggccaggaaccgtaaaaaggccgcgttgctggcgtttttccataggctccgcccccctgacgagcatcacaaaaatcgacgct
caagtcagaggtggcgaaacccgacaggactataaagataccaggcgtttccccctggaagctccctcgtgcgctctcctgttccga
ccctgccgcttaccggatacctgtccgcctttctcccttcgggaagcgtggcgctttctcatagctcacgctgtaggtatctcagttcggtgt
aggtcgttcgctccaagctgggctgtgtgcacgaaccccccgttcagcccgaccgctgcgccttatccggtaactatcgtcttgagtcca
acccggtaagacacgacttatcgccactggcagcagccactggtaacaggattagcagagcgaggtatgtaggcggtgctacaga

gttcttgaagtggtggcctaactacggctacactagaagaacagtatttggtatctgcgctctgctgaagccagttaccttcggaaaaag
agttggtagctcttgatccggcaaacaaaccaccgctggtagcggtggtttttttgtttgcaagcagcagattacgcgcagaaaaaaag
gatctcaagaagatcctttgatcttttctacggggtctgacgctcagtggaacgaaaactcacgttaagggattttggtcatgagattatca
aaaaggatcttcacctagatccttttaaattaaaaatgaagttttaaatcaatctaaagtatatatgagtaaacttggtctgacagttacca
atgcttaatcagtgaggcacctatctcagcgatctgtctatttcgttcatccatagttgcctgactccccgtcgtgtagataactacgatacg
ggagggcttaccatctggccccagtgctgcaatgataccgcgagacccacgctcaccggctccagatttatcagcaataaaccagcc
agccggaagggccgagcgcagaagtggtcctgcaactttatccgcctccatccagtctattaattgttgccgggaagctagagtaagt
agttcgccagttaatagtttgcgcaacgttgttgccattgctacaggcatcgtggtgtcacgctcgtcgtttggtatggcttcattcagctccg
gttcccaacgatcaaggcgagttacatgatcccccatgttgtgcaaaaaagcggttagctccttcggtcctccgatcgttgtcagaagta
agttggccgcagtgttatcactcatggttatggcagcactgcataattctcttactgtcatgccatccgtaagatgcttttctgtgactggtga
gtactcaaccaagtcattctgagaatagtgtatgcggcgaccgagttgctcttgcccggcgtcaatacgggataataccgcgccacat
agcagaactttaaaagtgctcatcattggaaaacgttcttcggggcgaaaactctcaaggatcttaccgctgttgagatccagttcgatg
taacccactcgtgcacccaactgatcttcagcatcttttactttcaccagcgtttctgggtgagcaaaaacaggaaggcaaaatgccgc
aaaaaagggaataagggcgacacggaaatgttgaatactcatactcttcctttttcaatattattgaagcatttatcagggttattgtctcat
gagcggatacatatttgaatgtatttagaaaaataaacaaataggggttccgcgcacatttccccgaaaagtgccacctgacggatcg
cttgcctgtaacttacacgcgcctcgtatcttttaatgatggaataatttgggaatttactctgtgtttatttatttttatgttttgtatttggattttaga
aagtaaataaagaaggtagaagagttacggaatgaagaaaaaaaaataaacaaaggtttaaaaaatttcaacaaaaagcgtactt
tacatatatatttattagacaagaaaagcagattaaatagatatacattcgattaacgataagtaaaatgtaaaatcacaggattttcgtg
tgtggtcttctacacagacaagatgaaacaattcggcattaatacctgagagcaggaagagcaagataaaaggtagtatttgttggcg
atccccctagagtcttttacatcttcggaaaacaaaaactatttttttctttaatttctttttttactttctatttttaatttatatatttatattaaaaaattt
aaattataattatttttatagcacgtgatgaaaaggaccctaagaaaccattattatcatgacattaacctataaaaataggcgtatcacg
aggccctttcgtctcgcgcgtttcggtgatgacggtgaaaacctctgacacatgcagctcccggagacggtcacagcttgtctgtaagc
ggatgccgggagcagacaagcccgtcagggcgcgtcagcgggtgttggcgggtgtcggggctggcttaactatgcggcatcagag
cagattgtactgagagtgcaccataaattcccgttttaagagcttggtgagcgctaggagtcactgccaggtatcgtttgaacacggcat
tagtcagggaagtcataacacagtcctttcccgcaattttctttttctattactcttggcctcctctagtacactctatatttttttatgcctcggtaa
tgattttcattttttttttccacctagcggatgactcttttttttttcttagcgattggcattatcacataatgaattatacattatataaagtaatgtgat
ttcttcgaagaatatactaaaaaatgagcaggcaagataaacgaaggcaaagatgacagagcagaaagccctagtaaagcgtatt
acaaatgaaaccaagattcagattgcgatctctttaaagggtggtcccctagcgatagagcactcgatcttcccagaaaaagaggca
gaagcagtagcagaacaggccacacaatcgcaagtgattaacgtccacacaggtatagggtttctggaccatatgatacatgctctg
gccaagcattccggctggtcgctaatcgttgagtgcattggtgacttacacatagacgaccatcacaccactgaagactgcgggattg
ctctcggtcaagcttttaaagaggccctaggggccgtgcgtggagtaaaaaggtttggatcaggatttgcgcctttggatgaggcacttt
ccagagcggtggtagatctttcgaacaggccgtacgcagttgtcgaacttggtttgcaaagggagaaagtaggagatctctcttgcga
gatgatcccgcattttcttgaaagctttgcagaggctagcagaattaccctccacgttgattgtctgcgaggcaagaatgatcatcaccgt
agtgagagtgcgttcaaggctcttgcggttgccataagagaagccacctcgcccaatggtaccaacgatgttccctccaccaaaggtg
ttcttatgtagtgacaccgattatttaaagctgcagcatacgatatatata

Plasmid pAF24 (pCTR1-VC83_00191-yeGFP) tCTR1 (Addgene plasmid 2223172)

*Legend*

CTR1 Promoter UNDERLINED

VC83_00191

Dual Streptag linker

GFP

CTR1 Terminator **BOLD**

*Plasmid Sequence:*

catgtgtatatatgtatacctatgaatgtcagtaagtatgtatacgaacagtatgatactgaagatgacaaggtaatgcatcattctatacg
tgtcattctgaacgaggcgcgctttcctttttttctttttgctttttcttttttttctcttgaactcgacggatctatgcggtgtgaaataccgcacaga
tgcgtaaggagaaaataccgcatcaggaaattgtaagcgttaatattttgttaaaattcgcgttaaatttttgttaaatcagctcattttttaac
caataggccgaaatcggcaaaatcccttataaatcaaaagaatagaccgagatagggttgagtgttgttccagtttggaacaagagtc
cactattaaagaacgtggactccaacgtcaaagggcgaaaaaccgtctatcagggcgatggcccactacgtgaaccatcaccctaa
tcaagttttttggggtcgaggtgccgtaaagcactaaatcggaaccctaaagggagcccccgatttagagcttgacggggaaagccg
gcgaacgtggcgagaaaggaagggaagaaagcgaaaggagcgggcgctagggcgctggcaagtgtagcggtcacgctgcgc
gtaaccaccacacccgccgcgcttaatgcgccgctacagggcgcgtccattcgccattcaggctgcgcaactgttgggaagggcgat
cggtgcgggcctcttcgctattacgccagctggcgaaagggggatgtgctgcaaggcgattaagttgggtaacgccagggttttccca
gtcacgacgttgtaaaacgacggccagtgaattgtaatacgactcactatagggcgaattggagctccaccgcggtggcggccgctc
tagaactagtcacagacattaacccacagtacagacactgcgacaacgtggcaattcgtcgcaatacaacgacacaagtccgattg
ttcctcttcaggagcttcctgaaccaaactttttccgcaaggccgcattttgaaccgtattttgctcgttccagcctttccacgtttttgttatcta
agcaacttggcacatttccctactatactacaaaccgatacgtaaatacttccctaaatagcatatgaattattcagtaatttttaaggatcg
aaactgcacctcaactattcgttactgtggttatgttctcatgtattgatgcaaatcatgggatatttgctcaagacgacggtaaaatgagc
aaaaatggcacgatcctgaaaagagcacttttcaagattcgggctacaaaatgcaacataaaaaatgttgtattgtcatctcgagagg
gtcttgtatgttttattcctcttatgattagttcacattagtaaaacagatacgcagtgtgctcttaataaacaactactccatagctttatttgca
taacaaaacttttaagcacaaacttaaacaggtggagtaatagttcggcggcgactcaaattacatttgttggaagaatcgaatagaa
aataaaaaaaagtgtattatatttgacattcaaatatggctgataatccattcgcaacatcaacgcctacttctggtgatgatatgtctggc
atggatatgtctcatggttcatcacacggcagttctcatgggtcatcgtccggcatgtctatggtcatgacctttcagaataatcctagcac
gcctttgttctctacagcgtggactcctaccggtacggggtcgtatgctgggacttgcattttttttaatcgtcttcgctgttctgtttagggttttatt
ggctcttaaagcgcgtcaagaagcaagatggcttgattgcgaaatgcatagaagatacgtggcggtcgctggcaagccaggcttaa
gagaacgtgttgcactacacaaggacgcgaaagcagtcgttttatctgaaaatggcgttgaggaagaggtagtggtggttcaaagaa
agggagaaatgacctcgccatggagggtttccgttgatcctcttagagcagtggtagatactgtgatagctgggatgggttacttgttaat
gcttgcggttatgactatgaacgtggggtacttcctttcagtcctagcaggagtgttcttgggtagtttagccattggtagatacactacttca
tacgaaggacacggtgggtccggcggttcatggtcacacccccaaccagaaaaaggtggtggatcaggagggtcttggagtcacc
ctcaacctgaaaagggaggatcctcgaaaggtgaagaattattcactggtgttgtcccaattttggttgaattagatggtgatgttaatggt
cacaaattttctgtctccggtgaaggtgaaggtgatgctacttacggtaaattgaccttaaaatttatttgtactactggtaaattgccagttc
catggccaaccttagtcactactttcggttatggtgttcaatgttttgcgagatacccagatcacatgaaacaacatgactttttcaagtctg
ccatgccagaaggttatgttcaagaaagaactatttttttcaaagatgacggtaactacaagaccagagctgaagtcaagtttgaaggt
gataccttagttaatagaatcgaattaaaaggtattgattttaaagaagatggtaacattttaggtcacaaattggaatacaactataactc
tcacaatgtttacatcatggctgacaaacaaaagaatggtatcaaagttaacttcaaaattagacacaacattgaagatggttctgttca
attagctgaccattatcaacaaaatactccaattggtgatggtccagtcttgttaccagacaaccattacttatccactcaatctgccttatc
caaagatccaaacgaaaagacagaccacatggtcttgttagaatttgttactgctgctggtattacccatggtatggatgaattgtacaa
aggatc**ctaataactagacaaggttcgtccacctatatcttcttcttccaatatctctatacatcaaaataagaatatcgttcatttc**
**tcagaagctcaaaaaaaatctcaaaatggatacttctaatgccctcatgaagaggaaagaaaataatagacacagcgacct**
**tcctataaataacagacttgtatatagttgaatgtctaagtaattttaattcaaaaatccttaactatattaccaccttgcaataata**
**agagtgaagtgttaaaaatgatacttgcttgccttcatcgagttatcgttgtctcccacgtttcttttttctcgttggacagaattgc**
**ttccgcgcgaacattttccacttttatttgaaagaggacattattcagatcgtgtcagtctcatctaattggcaaagtggcaatttt**
**cgaaataacaagataactgcataaagtaacacttgtcgaattgaaagtattttgccagtgatatttaggttcgagtaaagaaaa**
**tttcataaagaaatcaacaagacacaatgctg**gaaatctgctcgtcagtggtgctcagaattcgatatcaagcttatcgataccgtcg

acctcgagggggggcccggtaccagcttttgttccctttagtgagggttaatttcgagcttggcgtaatcatggtcatagctgtttcctgtgtg
aaattgttatccgctcacaattccacacaacatacgagccggaagcataaagtgtaaagcctggggtgcctaatgagtgagctaactc
acattaattgcgttgcgctcactgcccgctttccagtcgggaaacctgtcgtgccagctgcattaatgaatcggccaacgcgcggggag
aggcggtttgcgtattgggcgctcttccgcttcctcgctcactgactcgctgcgctcggtcgttcggctgcggcgagcggtatcagctcact
caaaggcggtaatacggttatccacagaatcaggggataacgcaggaaagaacatgtgagcaaaaggccagcaaaaggccag
gaaccgtaaaaaggccgcgttgctggcgtttttccataggctccgcccccctgacgagcatcacaaaaatcgacgctcaagtcagag
gtggcgaaacccgacaggactataaagataccaggcgtttccccctggaagctccctcgtgcgctctcctgttccgaccctgccgctta
ccggatacctgtccgcctttctcccttcgggaagcgtggcgctttctcatagctcacgctgtaggtatctcagttcggtgtaggtcgttcgct
ccaagctgggctgtgtgcacgaaccccccgttcagcccgaccgctgcgccttatccggtaactatcgtcttgagtccaacccggtaag
acacgacttatcgccactggcagcagccactggtaacaggattagcagagcgaggtatgtaggcggtgctacagagttcttgaagtg
gtggcctaactacggctacactagaagaacagtatttggtatctgcgctctgctgaagccagttaccttcggaaaaagagttggtagctc
ttgatccggcaaacaaaccaccgctggtagcggtggtttttttgtttgcaagcagcagattacgcgcagaaaaaaaggatctcaagaa
gatcctttgatcttttctacggggtctgacgctcagtggaacgaaaactcacgttaagggattttggtcatgagattatcaaaaaggatctt
cacctagatccttttaaattaaaaatgaagttttaaatcaatctaaagtatatatgagtaaacttggtctgacagttaccaatgcttaatcag
tgaggcacctatctcagcgatctgtctatttcgttcatccatagttgcctgactccccgtcgtgtagataactacgatacgggagggcttac
catctggccccagtgctgcaatgataccgcgagacccacgctcaccggctccagatttatcagcaataaaccagccagccggaagg
gccgagcgcagaagtggtcctgcaactttatccgcctccatccagtctattaattgttgccgggaagctagagtaagtagttcgccagtt
aatagtttgcgcaacgttgttgccattgctacaggcatcgtggtgtcacgctcgtcgtttggtatggcttcattcagctccggttcccaacga
tcaaggcgagttacatgatcccccatgttgtgcaaaaaagcggttagctccttcggtcctccgatcgttgtcagaagtaagttggccgca
gtgttatcactcatggttatggcagcactgcataattctcttactgtcatgccatccgtaagatgcttttctgtgactggtgagtactcaacca
agtcattctgagaatagtgtatgcggcgaccgagttgctcttgcccggcgtcaatacgggataataccgcgccacatagcagaacttta
aaagtgctcatcattggaaaacgttcttcggggcgaaaactctcaaggatcttaccgctgttgagatccagttcgatgtaacccactcgt
gcacccaactgatcttcagcatcttttactttcaccagcgtttctgggtgagcaaaaacaggaaggcaaaatgccgcaaaaaaggga
ataagggcgacacggaaatgttgaatactcatactcttcctttttcaatattattgaagcatttatcagggttattgtctcatgagcggataca
tatttgaatgtatttagaaaaataaacaaataggggttccgcgcacatttccccgaaaagtgccacctgacggatcgcttgcctgtaactt
acacgcgcctcgtatcttttaatgatggaataatttgggaatttactctgtgtttatttatttttatgttttgtatttggattttagaaagtaaataa
gaaggtagaagagttacggaatgaagaaaaaaaaaataaacaaaggtttaaaaaatttcaacaaaaagcgtactttacatatatattt
attagacaagaaaagcagattaaatagatatacattcgattaacgataagtaaaatgtaaaatcacaggattttcgtgtgtggtcttctac
acagacaagatgaaacaattcggcattaatacctgagagcaggaagagcaagataaaaggtagtatttgttggcgatccccctaga
gtcttttacatcttcggaaaacaaaaactattttttctttaatttctttttttactttctatttttaatttatatatttatattaaaaaatttaaattataattat
ttttatagcacgtgatgaaaaggaccctaagaaaccattattatcatgacattaacctataaaaataggcgtatcacgaggccctttcgt
ctcgcgcgtttcggtgatgacggtgaaaacctctgacacatgcagctcccggagacggtcacagcttgtctgtaagcggatgccggg
agcagacaagcccgtcagggcgcgtcagcgggtgttggcgggtgtcggggctggcttaactatgcggcatcagagcagattgtactg
agagtgcaccataaattcccgttttaagagcttggtgagcgctaggagtcactgccaggtatcgtttgaacacggcattagtcagggaa
gtcataacacagtcctttcccgcaattttctttttctattactcttggcctcctctagtacactctatatttttttatgcctcggtaatgattttcattttttt
tttccacctagcggatgactcttttttttttcttagcgattggcattatcacataatgaattatacattatataaagtaatgtgatttcttcgaagaat
atactaaaaaatgagcaggcaagataaacgaaggcaaagatgacagagcagaaagccctagtaaagcgtattacaaatgaaac
caagattcagattgcgatctctttaaagggtggtcccctagcgatagagcactcgatcttcccagaaaaagaggcagaagcagtagc
agaacaggccacacaatcgcaagtgattaacgtccacacaggtatagggtttctggaccatatgatacatgctctggccaagcattcc
ggctggtcgctaatcgttgagtgcattggtgacttacacatagacgaccatcacaccactgaagactgcgggattgctctcggtcaagc
ttttaaagaggccctaggggccgtgcgtggagtaaaaaggtttggatcaggatttgcgcctttggatgaggcactttccagagcggtggt
agatctttcgaacaggccgtacgcagttgtcgaacttggtttgcaaagggagaaagtaggagatctctcttgcgagatgatcccgcattt
tcttgaaagctttgcagaggctagcagaattaccctccacgttgattgtctgcgaggcaagaatgatcatcaccgtagtgagagtgcgtt
caaggctcttgcggttgccataagagaagccacctcgcccaatggtaccaacgatgttccctccaccaaaggtgttcttatgtagtgac
accgattatttaaagctgcagcatacgatatatata

Recombinant protein expressed by pAF24

*Legend*

- *Pd*CTR1a (VC83_00191)
- Strep-tag II Linker
- yeGFP

*Protein Sequence:*

MADNPFATSTPTSGDDMSGMDMSHGSSHGSSHGSSSGMSMVMTFQNNPSTPLFSTAWTPT
GTGSYAGTCIFLIVFAVLFRVLLALKARQEARWLDCEMHRRYVAVAGKPGLRERVALHKDAKAV
VLSENGVEEEVVVVQRKGEMTSPWRVSVDPLRAVVDTVIAGMGYLLMLAVMTMNVGYFLSVL
AGVFLGSLAIGRYTTSYEGHGGSGGSWSHPQPEKGGGSGGSWSHPQPEKGGSSKGEELFT
GVVPILVELDGDVNGHKFSVSGEGEGDATYGKLTLKFICTTGKLPVPWPTLVTTFGYGVQCFAR
YPDHMKQHDFFKSAMPEGYVQERTIFFKDDGNYKTRAEVKFEGDTLVNRIELKGIDFKEDGNIL
GHKLEYNYNSHNVYIMADKQKNGIKVNFKIRHNIEDGSVQLADHYQQNTPIGDGPVLLPDNHYL
STQSALSKDPNEKTDHMVLLEFVTAAGITHGMDELYKGS*
