## Supplemental Table 1 for "Characterization of a High-Affinity Copper Transporter in the White-Nose Syndrome Causing Fungal Pathogen *Pseudogymnoascus destructans*"

**SI Table 1.** Table of primers used in this study.

| Component | Template | Primers | Primer Sequence |
|---|---|---|---|
| CTR Promoter – GFP terminator cassette | Pytk110-DN6 | oRLP497 | cataggtctcactagtcacagacattaacccacagtacagacac |
| | | oRLP498 | ctaacggtctcgaattctgagcaccactgacgagcagatt |
| RT-qPCR | Actin cDNA | oRLP478 | ATCACACCTTCTACAACGAGC |
| | | oRLP479 | GGCGTTGAAAGTCTCGAAAAC |
| RT-qPCR | *Pd*Ctr1a cDNA | oRLP198 | GACAAGTCCGTGGAGAGTTAG |
| | | oRLP199 | AAGCTACCTAGGAAAACGCC |
| RT-qPCR | *Pd*Ctr1b cDNA | oRLP200 | CCCCATAACACTCACCATCTC |
| | | oRLP201 | GTCTCAGCAAGTTCCTCGTG |
